# *CrHCF244* is required for *psbA* translation in *Chlamydomonas reinhardtii*

**DOI:** 10.1101/2024.05.30.596694

**Authors:** Xiaozhuo Wang, Guannan Wang, Lexi A. Cheramie, Cuiping Zhao, Maheshi Dassanayake, James V. Moroney, David J. Vinyard

## Abstract

Translation of *psbA*, the chloroplast gene that encodes the D1 subunit of Photosystem II (PSII), is important for both PSII biogenesis and repair. The translation of the *psbA* transcript in the chloroplast is under the control of nuclear gene products. Using a *Chlamydomonas* forward genetic screen and whole genome sequencing, we found a mutant defective in PSII activity and mapped the causative gene to be the homolog of *Arabidopsis High Fluorescence* (*HCF244*) gene, or *CrHCF244*. We then demonstrated that CrHCF244 is required for *psbA* translation in the alga, consistent with the function of HCF244 in *Arabidopsis*. The *Arabidopsis HCF244* gene also partially complemented the algal mutant. These results experimentally support the functional conservation of the homologs in green algae and land plants. Intriguingly, the *CrHCF244* mutant also exhibited a relatively high rate of suppressor mutants, pointing to the presence of alternative factor(s)/pathway(s) for D1 translation control. The establishment of *CrHCF244* as a *psbA* translation factor in *Chlamydomonas* showed the similarities in *psbA* translation regulation in algae and plants. The future identification of the alternative factor(s) in this alga will provide insights on *psbA* translation in plants.

**Highlight:** We identified CrHCF244 as a translation factor of *psbA* in *Chlamydomonas*. *Arabidopsis HCF244* partially complements *Chlamydomonas* Δ*CrHCF244* mutant, indicating semi-conservation of the function of this gene between organisms. Suppressor mutants of Δ*CrHCF244* suggest the presence of alternative translation factors in *psbA* translation.

## Introduction

Photosystem II (PSII) is a water-plastoquinone oxidoreductase that enables the capture and conversion of light energy into storable chemical energy. PSII is the first component in the photosynthetic electron transport chain that generates ATP and NADPH for CO_2_ fixation. Though much is known about the subunit composition and structure of PSII (Shen, 2015), its biogenesis is less well understood. As a large protein complex containing over 20 subunits encoded by both nuclear and chloroplast genomes and containing various cofactors, the biogenesis of PSII is complicated (Lu, 2016; Nickelsen and Rengstl, 2013; Nixon *et al*., 2010). PSII biogenesis is a stepwise and highly coordinated process involving the synthesis of subunits, integration into the membrane, assembly of protein subunits, and association of cofactors (Lu, 2016; Nickelsen and Rengstl, 2013; Nixon *et al*., 2010). PSII is also subject to photoinhibition and a repair process is needed to restore its activity, during which the D1 subunit needs to be degraded and a new copy of the subunit synthesized *de novo* (Järvi *et al*., 2015).

Translation of chloroplast genes encoding subunits of photosynthetic complexes including PSII have been found to be widely regulated by nuclear gene products, which are termed as regulators of organelle genes (ROGEs). As the control of gene expression in chloroplasts occurs prevalently at post-transcriptional levels, including mRNA stabilization and/or its translation, ROGEs are categorized as M and T factors, which refer to protein factors involved in maturation or stabilization of the target mRNAs and translation regulation of the mRNAs, respectively. Genetic and biochemical approaches have identified many ROGEs in regulating PSII in *Chlamydomonas* (e.g. Tba1 (Somanchi *et al*., 2005), RBP63/DLA2 (Bohne *et al*., 2013; Ossenbuhl *et al*., 2002), RBP40/RB38 (Schwarz *et al*., 2007), MBB1 (Vaistij *et al*., 2000), MBC1 (Cavaiuolo *et al*., 2017), TBC2 (Auchincloss *et al*., 2002)), PSI (e.g. TAA1 (Lefebvre-Legendre *et al*., 2015)), Cyt b_6_f (e.g. MCG1 (Wang *et al*., 2015)), and ATP synthase (e.g. TDA1 (Drapier *et al*., 1992; Eberhard *et al*., 2011), MDA1 (Viola *et al*., 2019)). Recently a systematic screening and characterization discovered several gene specific ROGEs and master regulators responsible for regulating the expression of multiple ROGEs to be involved in biogenesis of photosynthetic machinery (Kafri *et al*., 2023).

Among PSII genes, the translation of *psbA* encoding the D1 subunit is important for both PSII *de novo* biogenesis and repair. *psbA* gene expression control has been extensively studied in cyanobacteria, algae, and plants (Mulo *et al*., 2012). A few protein factors have been identified as involved in *psbA* translation in *Chlamydomonas*. Mayfield and coworkers biochemically identified *psbA* mRNA binding proteins RB47, RB60, RB38, and RB55, named based on their molecular size (Danon and Mayfield, 1991). The gene for RB47 was later cloned as a polyadenylate (poly (A)) binding protein, thus also named as chloroplast poly (A) binding protein cPAB1 (Yohn *et al*., 1998a; Yohn *et al*., 1998b). Though chloroplast mRNAs generally do not have poly (A) tails, there are A-rich regions in the UTRs, which could be potential binding sites. RB60 is a disulfide isomerase protein (Kim and Mayfield, 1997) which may regulate the redox state of RB47 and thus control its binding to *psbA* mRNA and the translation (Kim and Mayfield, 2002). Tba1, identified in a genetic screen, is a thioredoxin protein regulating *psbA* translation (Somanchi *et al*., 2005). Genetic screens have also found several *Chlamydomonas* nuclear mutants defective in *psbA* translation, some of which the causative genes are not mapped including F35 (Girard-Bascou *et al*., 1992; Yohn *et al*., 1996). Besides the dedicated protein factors involved in *psbA* translation, a “moonlighting” regulator was also identified. The dihydrolipoamide acetyltransferase subunit 2 (DLA2), a subunit of chloroplast pyruvate dehydrogenase complex (cpPDC) is involved in D1 translation regulation under mixotrophic growth condition (Bohne *et al*., 2013; Neusius *et al*., 2022; Ossenbuhl *et al*., 2002). DLA2 switches its role between acetate metabolism as a subunit of cpPDC, and regulator of D1 translation dissociating from the cpPDC complex and associating with thylakoid membrane through acetylation of its K197 residue (Neusius *et al*., 2022).

In land plants, two protein factors have been identified to be involved in *psbA* translation: High Chlorophyll Fluorescence proteins 173 and 244 (HCF173 and HCF244) (Chotewutmontri and Barkan, 2020; Link *et al*., 2012; Schult *et al*., 2007). Both are members of the short-chain dehydrogenase superfamily (Kavanagh *et al*., 2008; Moummou *et al*., 2012; Oppermann *et al*., 2001). HCF173 is required for *psbA* translation initiation and mRNA stability (Schult *et al*., 2007). HCF244, co-expressing with HCF173, is also involved in *psbA* translation, possibly working together with HCF173, as the double mutant has a more reduced plant size and fewer numbers of leaves (Link *et al*., 2012).

In this study, we used forward genetics to identify a PSII defective mutant in *Chlamydomonas* in which the causative gene was mapped to be a homolog of *Arabidopsis HCF244*. By examining the mutant, we provide evidence for its role in D1 translation in this alga. We show that the *Arabidopsis* homolog gene *HCF244* partially complements the *Chlamydomonas* mutant. Intriguingly, we observed a number of suppressor mutants of the *CrHCF244* mutant that showed partial recoveries of PSII activity. In an effort to examine the relationship of CrHCF244 to the previously identified translation activators RB60 and RB47 in *Chlamydomonas*, we generated the corresponding mutants using CRISPR-Cas9 but no effect on PSII activity was observed in those mutants. This study supports the functional conservation of the plant ortholog HCF244 in algae. Suppressor mutants suggest an alternative pathway of D1 translation control. Genetic studies add to our understanding of the components and their roles in D1 translation control.

## Materials and methods

### Strains and culture conditions

*Chlamydomonas* strain CC-4533 (Zhang *et al*., 2014) and *psbA* deletion strain Fud7 (Bennoun *et al*., 1986) were obtained from the *Chlamydomonas* Resource Center (CRC). CC-4533 is the background strain for the CLiP library. We refer to CC-4533 as WT here. The *ami6* mutant and the CRISPR-Cas9 knockout mutants (Δ*CrHCF244-T23*, Δ*RB47*, Δ*RB60-1, 2, 3*) were generated in the background of CC-4533. Cells were grown in Tris-acetate-phosphate (TAP) (Harris *et al*., 2009) or high salt (HS) (Sueoka, 1960) media with shaking at 25°C in an AlgaeTron incubator (Photon Systems Instruments). Growth tests were done by spotting cells onto TAP and/or HS plates with 1.5% (w/v) agar under white light at 60 µmol photons m^-2^ s^-1^ intensity.

### Chlamydomonas transformation

Electroporation was used in this study for the delivery of plasmid DNA (for generating random insertional mutants and complementation studies of mutants) and ribonucleoprotein (RNP) complexes (for generating CRISPR-Cas9 knockout mutants) into *Chlamydomonas* cells. Electroporation was done using MAX Efficiency^TM^ transformation reagent (Invitrogen) according to the manufacturer’s instructions. Briefly, cells from a 50 mL TAP culture with OD_750_ 0.3–0.5 were pelleted, washed twice with the transformation reagent and resuspended in 0.5–1 mL transformation reagent. For delivering plasmid, 110 µL of cell suspension was mixed with 1 µg plasmid DNA. For CRISPR Cas9 mutations, the 110 µL cell suspension was mixed with 10 µL Cas9-guide RNA (gRNA) RNP complex and 1 µg donor DNA. The mixture was added into a cuvette with 0.2 cm gap (Bio-Rad), and electroporated with Gene Pulser® II (Bio-Rad) at a voltage of 0.35 kV and capacitance of 25 µF. After electroporation, cells were kept at room temperature (RT) for 10 min, then transferred into 10 mL TAP with 40 mM sucrose and shaken in the dark overnight. Cells were then pelleted and spread onto selective plates. For complementation studies of *ami6* and for the Δ*CrHCF244-T23* mutant, plasmids C1 and C2 (see cloning construct) were electroporated into the mutants, and selected on HS plates.

### Whole genome sequencing and analysis

Genomic DNA was extracted from CC-4533 and *ami6* mutant using the phenol-chloroform-isoamyl method (Newman *et al*., 1990). The isolated DNA was further treated with RNase to remove RNA contamination and quantified using a Qubit fluorometer. Library preparation and Illumina sequencing was performed by Genewiz (NJ, USA). After quality check with FastQC (v0.11.7, http://www.bioinformatics.babraham.ac.uk/projects/fastqc/), paired-end reads were aligned to *Chlamydomonas reinhardtii* CC-4532 reference genome (v 5.6 and v6.1) (Merchant *et al*., 2007) from Phytozome V13 using bowtie2 (v2.3.5) (Langmead and Salzberg, 2012) with the default settings. Aligned reads were sorted and deduplicated using Picard (v2.25.2, https://broadinstitute.github.io/picard). We subsequently employed GRIDSS2 structural variant caller (v2.13.2) (Cameron *et al*., 2021) to identify rearrangement break points that generally represent structural variant events using the recommended workflow. Results were organized into VCF (Variant Call Format) files, and then filtered using StructuralVariantAnnotation (v1.18.0, DOI: 10.18129/B9.bioc.StructuralVariantAnnotation) in R. The resulting structural variants were further validated and supplemented with the ones identified by GATK (v4.1) (O’Connor and van der Auwera, 2020) to generate the final list of variants. Lastly, the identified variants for the WT and *ami6* mutant were compared to identify these that were only present in the mutant.

### Cloning constructs

Plasmid C1 was used for complementing *ami6* and CRISPR Cas9 knockout Δ*CrHCF244-T23* mutants with *CrHCF244* and the resulting transformants are *ami6-C1* and *T23-C1*. Plasmid C2 was used for complementing *ami6* and CRISPR Cas9 knockout Δ*CrHCF244-T23* mutants by *CrHCF244* with C-terminal tagged Venus and the resulting transformants are *ami6-C2* and *T23-C2*.

To construct the C1 plasmid, the open reading frame (ORF) of *CrHCF244* gene was amplified by Q5^®^ High-Fidelity DNA Polymerase (New England Biolabs) using CC-4533 genomic DNA as template; the linear vector was PCR amplified using pLM006 (Mackinder *et al*., 2016) as template with primers in Table S1 and the mCherry and His tag sequences were excluded. To construct C2, the above ORF of *CrHCF244* was used and the linear vector was PCR amplified using pLM005 (Mackinder *et al*., 2016) as template with primers in Table S1 (Venus sequence was included). The PCR products of the ORF of *CrHCF244* and linear vectors were assembled by Gibson assembly (Gibson et al., 2009) using Gibson Assembly^®^ Master Mix (New England Biolabs). For the vector construct to transform the *Arabidopsis HCF244* gene into *Chlamydomonas*, the coding sequence for the HCF244 protein without the transit peptide was codon optimized (IDT Codon Optimization Tool) and then fused to the sequence of the *Chlamydomonas* PsaD transit peptide on the 5’ end. The chimeric sequence was synthesized by Twist Bioscience. The synthesized fragment was used as template for PCR using primers in Table S1, after which it was assembled with the PCR product using primers in Table S1 from the backbone plasmid pLM006 (Mackinder *et al*., 2016), excluding mCherry and His tag sequence. Plasmids were verified by Sanger sequencing before transformation into *Chlamydomonas*.

### Generating CRISPR-Cas9 knockout mutants in Chlamydomonas

CRISPR-Cas9 was used for targeted mutagenesis to knock out the *CrHCF244*, *RB47* and *RB60* genes in this study. The gRNA design, *in vitro* assembly with Cas9 to form RNP and delivery of RNP and donor DNA to cells were performed according to a published method (Picariello *et al*., 2020) with minor modifications. Briefly, the crRNAs targeting the second exon of *CrHCF244* (Cre02.g142146), the fourth exon of *RB47* (Cre01.g039300) and the first exon of *RB60* (Cre02.g088200) were designed on the website of Integrated DNA Technologies (IDT). The lack of off-targeting was verified by Cas-OFFinder (Bae *et al*., 2014). The crRNAs and tracrRNA (sequences were shown in Table S1), synthesized by IDT, were first annealed and then assembled with Cas9 (Alt-R™ S.p. Cas9 Nuclease V3, IDT) at 37°C for 15 min. Donor DNA for editing each gene was PCR amplified with primers in Table S1 using the pKS-aphVIII-lox aphVIII plasmid (Heitzer and Zschoernig, 2007) or the pHyg3 plasmid (Berthold *et al*., 2002) as templates to generate antibiotic-resistance cassettes flanked by 40-bp homology arms of the Cas9 cleavage site. The RNP and donor DNA was electroporated into CC-4533 after removal of cell wall by autolysin. Transformants were selected on TAP plates with 20 µg/mL paromomycin or hygromycin. Colonies were screened with PCR using primers in Table S1 and/or Sanger sequencing.

### Spotting cells for growth tests

For spotting, TAP cultures with OD_750_ around 0.3–0.5 were used. Cells from 1 mL of culture were pelleted, washed with HS, and resuspended in 1 mL HS. OD_750_ was used to normalize the number of cells. Five µL cultures with serial dilutions were spotted onto TAP and HS plates. Plates were kept under 60 µmol photons m^-2^ s^-1^ at 25°C for growth.

### Chlorophyll fluorescence measurement

Chlorophyll a fluorescence F_v_/F_m_ ((F_m_-F_o_)/F_m_) was measured to indicate PSII activity (Genty *et al*., 1989). Cells in liquid cultures were measured with a double-modulation fluorometer (FL 6000, Photon Systems Instruments) after two minutes dark adaptation. Cells on plates were measured with a FluorCam (Photon Systems Instruments) after two minutes dark adaptation.

### Confocal microscopy for subcellular localization of CrHCF244

Confocal microscopy of *Chlamydomonas* cells was done according to (Mackinder *et al*., 2016), using a Leica SP8 microscope. The Venus tagged strain (*ami6-C2*) was grown in TAP to early exponential phase under light intensity of 60 µmol photons m^-2^ s^-1^. Around 8 µL of culture was mixed with an equal volume of 0.1% (w/v) low melting agarose on the slide and covered with a #1.5 cover slip. The observation was done using a 63x water immersion objective. White light laser at 512 nm was used for excitation of Venus fluorescence and emission between 520-540 nm was collected. Chlorophyll autofluorescence was excited at 561 nm and emission between 560-590 nm was collected as chlorophyll fluorescence.

### Sodium dodecyl sulfate-polyacrylamide electrophoresis (SDS-PAGE) and immunoblotting analysis

To analyze the abundance of photosynthetic proteins, *Chlamydomonas* cells were grown in liquid TAP under light intensity of 20 µmol photons m^-2^ s^-1^. The low light was used here to avoid possible secondary effects on the photosynthetic machinery in the mutants from high light intensity. Total protein was extracted using 2% SDS in 50 mM Tris-HCl (pH 8.0). A BCA assay was used to measure total protein concentration. Laemmli sample buffer was added to protein samples followed by incubation at room temperature for 10 min. Around 10 µg of protein was loaded to home-made 12% Tris-Glycine polyacrylamide gels containing 6 M urea. After electrophoresis, proteins were transferred to a PVDF membrane with pore size of 0.2 μm (Amersham™ Hybond™ P) through wet transfer in Tris-Glycine transfer buffer and blocked with 5% non-fat milk in Tris-buffered saline with 0.05% Tween 20 (TBST). Antibodies of PSII subunits D1 (Agrisera, AS05 084), D2 (Agrisera, AS06 146), CP43 (Agrisera, AS11 1787), CP47 (PhytoAB, PHY0319) and PsbO (Agrisera, AS06 142), and antibodies against PSI subunit PsaA (Agrisera, AS06 172) and Cyt b_6_f subunit Cyt f (a gift from Prof. Terry Bricker, LSU) were used. A rabbit polyclonal antibody against RB60 was made in this study using the RB60 peptide SEKMPPTIEFNQKNSDKIFN as the antigen by YenZym Antibodies LLC (https://www.yenzym.com). The PVDF membrane was incubated with diluted primary antibodies for 1 h at room temperature followed by washing and incubation with 1:20,000 diluted HRP-conjugated secondary antibody (Invitrogen, 31460) at room temperature for 1 h. ECL reagent (Bio-Rad, Clarity Western ECL Substrate) was applied to the membrane for chemiluminescence detection using a ChemiDoc XRS+ imaging system.

### Crude thylakoid membrane isolation and Blue-native (BN) PAGE

Crude thylakoid membranes from *Chlamydomonas* were prepared as described (Garcia-Cerdan *et al*., 2019) with minor modifications. Cells were resuspended in lysis buffer (25 mM HEPES-NaOH, pH 7.5, 10 mM NaCl, 5 mM MgCl_2_ and 5% [w/v] glycerol) and passed through a French press at gauge pressure 700 psi, 4°C to lyse cells. The lysate was centrifuged at 3000×*g*, 5 min, 4°C to remove unbroken cells and cell debris. The supernatant was then centrifuged at 30,000 × *g*, 30 min, 4°C. The pellets are used as crude thylakoid membranes. After resuspension in solubilization buffer (25 mM HEPES-NaOH, pH 7.5, 50 mM NaCl and 5% glycerol) at Chl concentration of 0.5 µg/µL, the thylakoid membranes were solubilized with 1% n-Dodecyl α-maltoside (α-DDM) (GLYCON Biochemicals GmbH) on ice for 20 min (Tokutsu *et al*., 2012), followed by centrifugation at 20,000 × *g* for 10 min at 4°C to remove unsolubilized thylakoids. The supernatant was run on a 5-12% BN-PAGE gel as previously described (Wang *et al*., 2016). Purified PSII from the *Chlamydomonas* B-His strain in which a His_6_ tag fused to the C-terminus of PsbB was used as a control (Suzuki *et al*., 2003).

### Chlorophyll measurement

Chlorophyll measurement was done using methanol extraction, as previously described (Porra *et al*., 1989). Cells were grown in TAP under light intensity of 35 µmol photons m^-2^ s^-1^ and in the dark, respectively.

### 14C-acetate pulse labeling

Pulse labeling of *Chlamydomonas* cells was done according to the literature (Calderon *et al*., 2023; Chua and Gillham, 1977; Spaniol *et al*., 2022). Briefly, 5 mL of TAP-grown cells under 20 µmol photons m^-2^ s^-1^ were pelleted and washed with HS. After resuspension in 0.5 mL HS, cells were incubated under 20 µmol photons m^-2^ s^-1^ with vigorous shaking for 1.5 h to deplete the cellular carbon pool. Then, 10 µg/mL cycloheximide was added to the culture to inhibit cytosolic translation followed by the addition of 20 µCi/mL ^14^C-acetic acid, sodium salt (Moravek, MC200) and incubated for 20 min under 20 µmol photons m^-2^ s^-1^. The culture was diluted into 10 volumes of cold TAP with 50 mM nonradioactive acetate and cells were pelleted and total protein was extracted. Protein samples were run on 12-18% Tris-Glycine SDS-PAGE gels with 6 M urea and then transferred from the gel to a PVDF membrane with pore size of 0.2 μm (Amersham™ Hybond™ P) (Amersham^TM^). The membrane was then exposed to a storage phosphor screen (Molecular Dynamics) for three weeks and imaged with a Typhoon^TM^ 8600 imager.

### Reverse transcription (RT)-PCR

Cell pellets from 10 mL TAP culture with OD750 of 0.3 were suspended in 1 mL TRIzol TM (Invitrogen). Total RNA was extracted using the Quick-RNA MiniPrep Kit (Zymo Research). ProtoScript® First Strand cDNA Synthesis Kit (New England Lab) was used to make cDNA using 500 ng RNA and an oligo-d(T) primer. No reverse transcriptase was added to the negative control group. Primers for *AtHCF244*(AtHCF244_1F, AtHCF244_2R), *RB47* (RB47_E4F and RB47_E7R), and *Actin* (Actin_F and Actin_R) are listed in Table S1.

## Results

### Screen and mapping of a mutant ami6 defective in PSII activity

To look for protein factors involved in PSII biogenesis, random insertional mutagenesis using a paromomycin cassette was used to generate a pool of mutants with resistance to this antibiotic. One mutant was identified which could not grow photoautotrophically and had no PSII activity, which we named *ami6* (Fig. 1A). To identify the candidate causative gene(s) in this mutant, we conducted whole genome sequencing. Analysis of the whole genome sequencing data using *Chlamydomonas reinhardtii* v6.1 (https://phytozome-next.jgi.doe.gov/info/CreinhardtiiCC_4532_v6_1) genome as the reference genome revealed a large deletion (∼347 kb) which mapped onto chromosome 15 (Fig. 1B). This genomic variance was verified by PCR and Sanger sequencing (primer sequences are in Supplementary Table S1 and results are in Supplementary Fig. S1). There are 28 genes (Supplementary Dataset S1) located in the deleted genome region. One of the deleted genes, Cre02.g142146, a homolog of *Arabidopsis HCF244*, was able to restore photoautotrophic growth and PSII activity in *ami6* (Fig. 1A). Adding the fluorescent protein Venus tag at the C-terminal of CrHCF244 did not affect the function of the protein (Fig. 1A and 1C). Of note, after we confirmed the causative gene in *ami6* in late May of 2022, we noticed an annotation on this gene in *Chlamydomonas* genome v6.1, the newly released version of the genome, that this gene is mutated in F35, a previously identified nuclear mutant defective in PSII (Girard-Bascou *et al*., 1992; Yohn *et al*., 1996) . However, there is no publication regarding on the mapping of the gene in F35. Our identification, mapping and complementation of *ami6* mutant verified the genetic relation of *CrHCF244* with PSII in *Chlamydomonas*.

**Figure 1.**
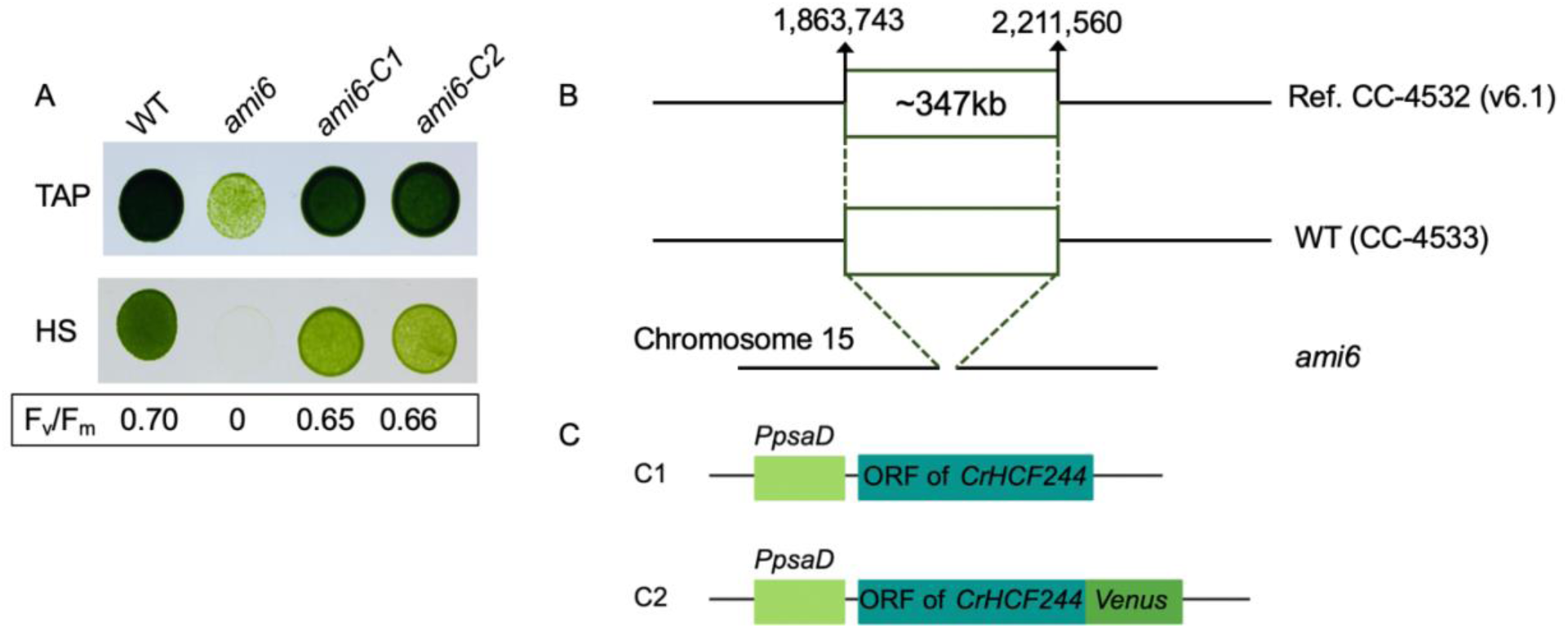
ami6 growth, PSII activity, and the mapping of genomic variation and the causative gene. **(A)** Growth of wild type (WT), *ami6* mutant and two complemented lines of *ami6*, under mixotrophic (TAP) and photoautotrophic (HS) conditions. *ami6-C1* is a representative transformant from transforming *ami6* with the C1 construct (illustrated in **(C)**); *ami6-C2* is a representative transformant from transforming *ami6* with the C2 construct (illustrated in **(C)**). F_v_/F_m_ values are shown for cultures grown on TAP. Spot test results and F_v_/F_m_ values are representative of two replicates. **(B)** Mapping of *ami6* genomic variation by whole genome sequencing revealed a ∼347 kb deletion on chromosome 15. **(C)** Illustration of genetic constructs for complementing *ami6* mutant. The C1 and C2 constructs both are under control of the *psaD* promoter (*PpsaD*). The constructs were used for transforming untagged CrHCF244 and Venus-tagged CrHCF244 into the *ami6* mutant.

### CRISPR-Cas9 to knock out CrHCF244

We next used CRISPR-Cas9 to generate a *CrHCF244* knockout mutant. The gRNA was designed to target the second exon of the gene, as illustrated in Fig. 2A. Based on PCR screening and Sanger sequencing (Supplementary Fig. S2A and Fig. S2B), transformant number 23 was selected, which is termed as Δ*CrHCF244-T23*. Δ*CrHCF244-T23* has a genomic variation at the Cas9 cleavage site in exon 2 of *CrHCF244*. For sequence alignment see Supplementary Fig. S2B. This mutation would result in the point mutation of F37 to G37 and a frame shift after the 37^th^ residue followed by a premature early stop codon, which abolished the function of the gene. Δ*CrHCF244-T23* could not grow photoautotrophically and has no PSII activity (Fig. 2B and C), consistent with the phenotype of *ami6*. Similar to *ami6*, *CrHCF244* restored the photosynthetic growth and PSII activity of Δ*CrHCF244-T23* (Fig. 2B). The complementation of both the *ami6* mutant and the CRISPR-Cas9 knockout mutant by *CrHCF244* verified that this gene is required for PSII function. The function is consistent with its homolog in plants (Chotewutmontri *et al*., 2020; Link *et al*., 2012).

**Figure 2.**
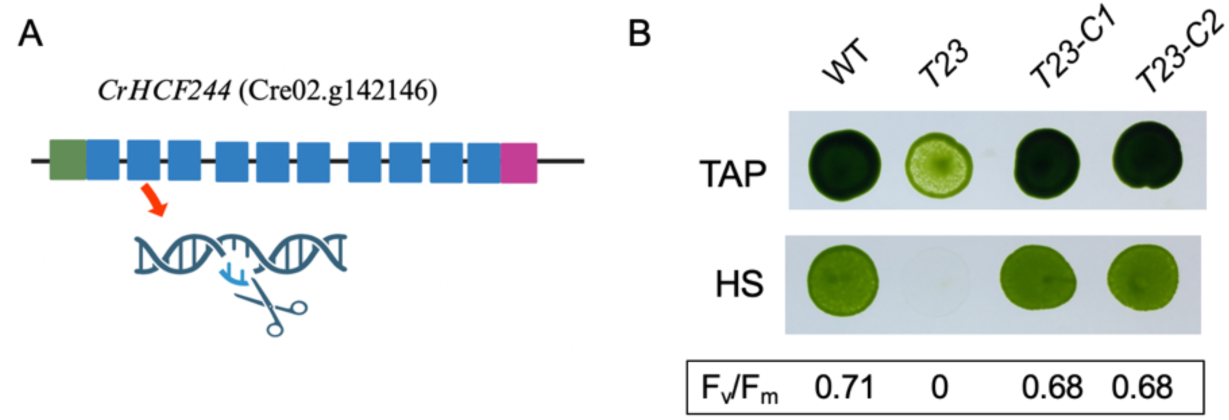
Growth and PSII activity of a CRISPR-Cas9 knockout CrHCF244 mutant. **(A)** *CrHCF244* gene map (5’ UTR as green box, exons as blue boxes, 3’ UTR as purple box) indicating the gRNA targeting site at exon 2 for CRISPR-Cas9 knockout/gene editing. **(B)** Growth of WT, CRISPR-Cas9 knockout *CrHCF244* mutant Δ*CrHCF244-T23* (shown as *T23* in **(B)** and **(C)**) and the two complemented strains (shown as *T23-C1* and *T23-C2* in **(B)** and **(C)**), which are representative transformants from transforming Δ*CrHCF244-T23* with C1 and C2 construct (illustrated in Fig. 1C), respectively, under mixotrophic (TAP) and photoautotrophic (HS) conditions. **(C)** F_v_/F_m_ values for strains grown on TAP. Spot test results and F_v_/F_m_ values are representative of two replicates.

### Mutation of CrHCF244 specifically affects accumulation of PSII among photosynthetic complexes

To test the accumulation of photosynthetic complexes in the absence of CrHCF244, immunoblotting of subunits of PSII, PSI and Cyt b_6_f was performed. As shown in Fig. 3A, PSII core subunits including D1, D2, CP43, CP47 were drastically reduced in the *ami6* mutant and Δ*CrHCF244-T23* mutant, whereas the abundance of subunits of PSI and Cyt b_6_f did not change in the mutants. The extrinsic subunit PsbO also showed a decrease in abundance, though to a lesser level than those core subunits. Introducing *CrHCF244* into the two mutants restored the accumulation of PSII subunits, as shown in the complemented lines (Fig. 3A). With respect of the PSII complexes, immunoblotting of the protein complexes separated on BN-PAGE showed that none of the PSII complexes accumulated to a significant level in *ami6* compared to WT (Fig. 3B and C). In the complemented strains, PSII complexes were restored, as shown in Fig. S3. The results showed absence of CrHCF244 impaired both accumulation of individual subunits and assembly of PSII complexes.

**Figure 3.**
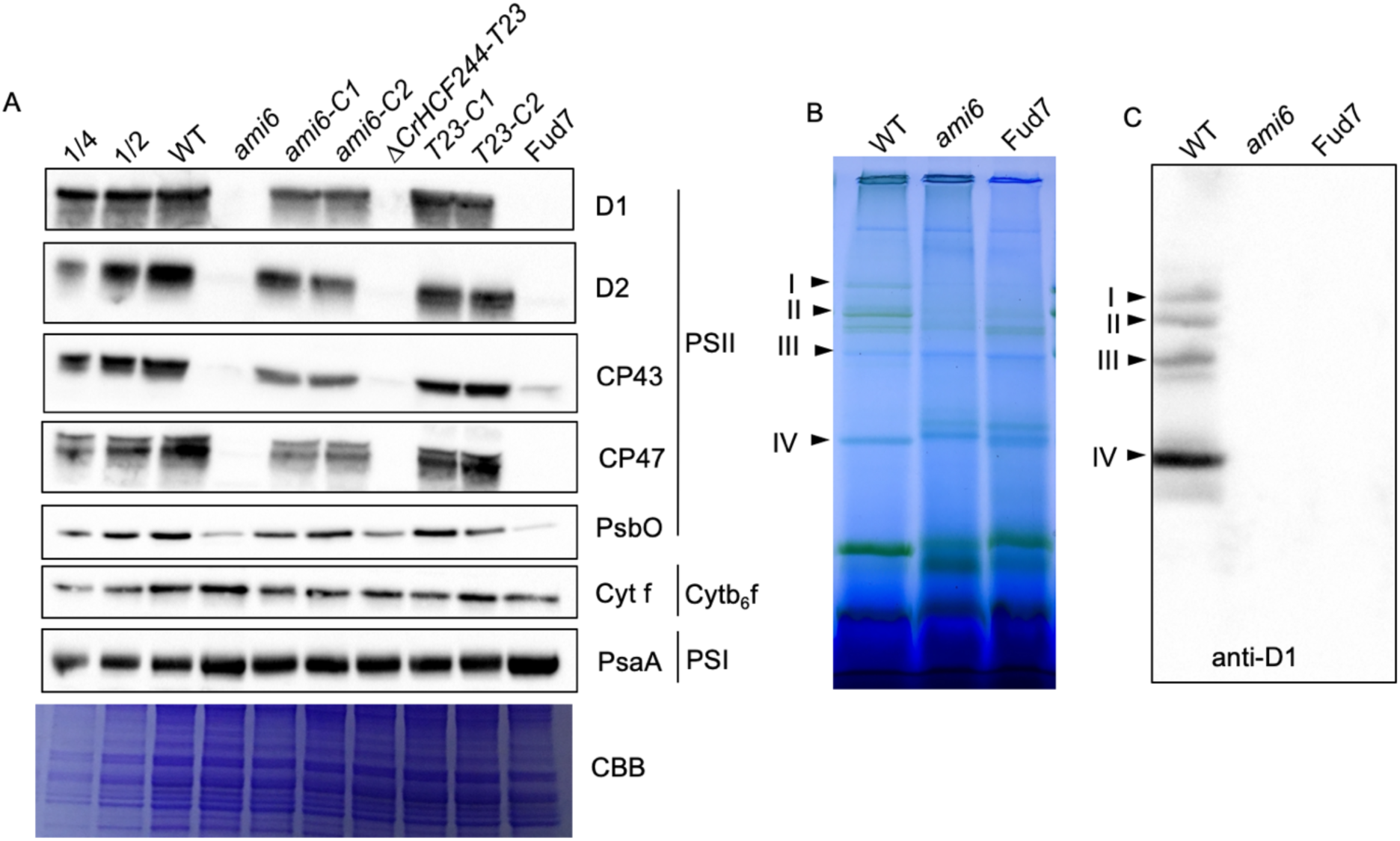
Accumulation of photosynthetic protein complexes. **(A)** Immunoblots of photosynthetic proteins in WT, *ami6* and its two complemented lines (*ami6-C1*, *ami6-C2*), Δ*CrHCF244-T23* and its two complemented lines (*T23-C1*, *T23-C2*) and Fud7 (*psbA* deletion mutant). Coomassie brilliant blue (CBB) stain was shown as a loading control. Data are representative of two biological replicates. **(B)** Blue-Native PAGE (BN-PAGE) separation of photosynthetic protein complexes. Arrows indicate the positions of different PSII complexes. **(C)** Immunoblotting analysis using D1 antibody of the PSII complexes. The immunoblotting was performed after transferring the protein complexes on the BN-PAGE to PVDF membrane. Arrows indicate PSII complexes, I and II: PSII super-complexes, III: PSII dimer, IV: PSII monomer.

### Mutation of CrHCF244 appears to abolish psbA translation

We used ^14^C-acetate pulse labeling to look at the synthesis of PSII subunits in *ami6* and the CRISPR-Cas9 mutants. Fig. 4A showed that in the mutants the synthesis of D1 subunit is not observable, whereas the synthesis of D2 and CP43 occurred similarly to WT. The synthesis of CP47 was absent in the mutants. This is due to the assembly-controlled translation which requires the presence of D1 for translation of CP47 (Minai *et al*., 2006). The transcription of *psbA* gene is slightly decreased in the mutant (Fig. 4B). This indicates that the decrease of D1 synthesis is not caused by defects in transcription but due to defects in translation.

**Figure 4.**
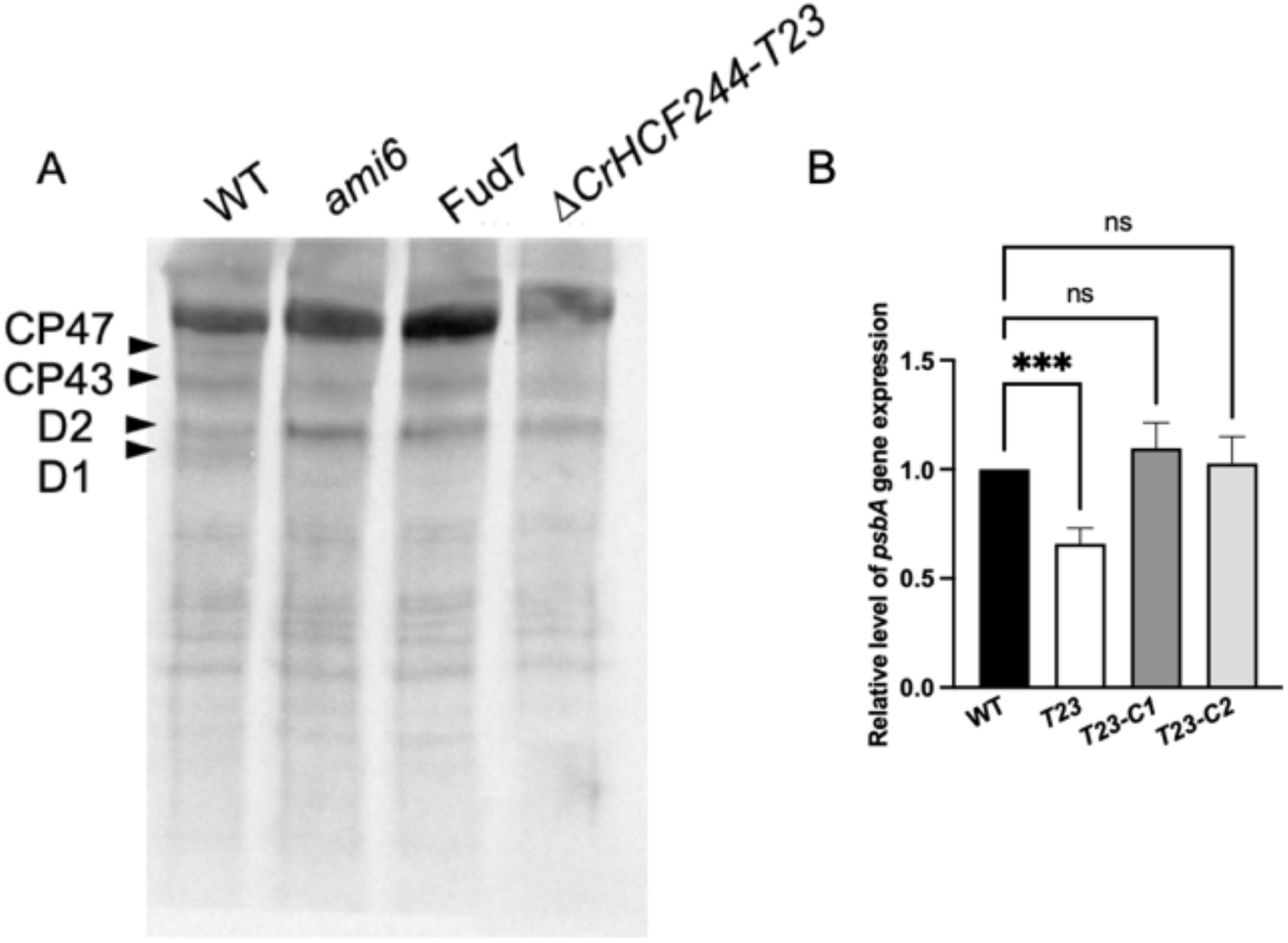
D1 synthesis and psbA transcription. **(A)** ^14^C-acetate pulse-labeling of WT, *ami6* mutant, Fud7 (*psbA* deletion) and CRISPR-Cas9 knockout mutant Δ*CrHCF244-T23*. Cells were labeled for 20 min in the presence of 10 µg/mL cycloheximide to inhibit cytosol translation. PSII major subunits are shown by the arrows. Data are representative of two biological replicates. **(B)** Quantitative real time PCR (qRT-PCR) to detect *psbA* transcription change in WT, Δ*CrHCF244-T23* and its two complemented lines (*T23-C1*, *T23-C2*). Data are presented as means ± Standard Deviation (n = 4). *** indicates p<0.001 and ns indicates non-significant with One-way ANOVA. The sequences of primers are shown in Table S1.

### Arabidopsis HCF244 complements the ami6 mutant

To test whether the *Arabidopsis* homolog gene could complement the *Chlamydomonas* mutant, we made a construct in which the sequence of the transit peptide of PsaD from *Chlamydomonas* (CrPsaD) was fused with the sequence of the mature *Arabidopsis* HCF244 protein (Fig. 5A). The construct was then transformed into *ami6* and transformants were selected under photoautotrophic conditions. The results (Fig. 5B) showed that *AtHCF244* can partially complement the *Chlamydomonas* mutant. Transgene expression was verified at the mRNA level by reverse transcription (RT)-PCR, as shown in Fig. 5C.

**Figure 5.**
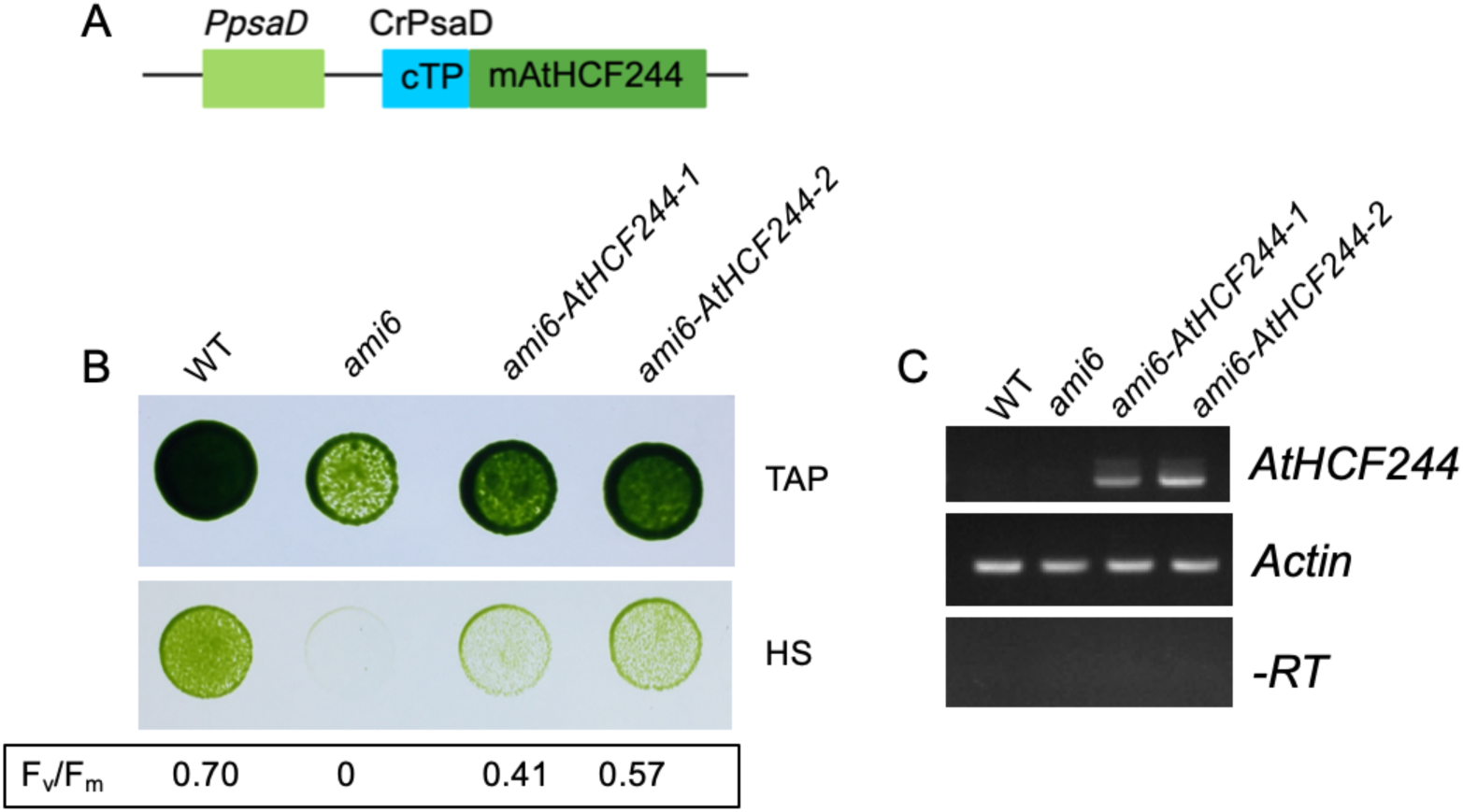
Complementation of Chlamydomonas ami6 mutant with the Arabidopsis HCF244 gene. **(A)** Schematic of the construct for transforming *ami6* with the *Arabidopsis HCF244* gene. Mature AtHCF244 protein (mAtHCF244) in fusion with a chloroplast transit peptide (cTP) derived from PsaD in *Chlamydomonas* was expressed under control of *psaD* promoter (*PpsaD*) **(B)** Growth and PSII activity of WT, *ami6* mutant, and the two complemented strains under mixotrophic (TAP) and photoautotrophic (HS) conditions. F_v_/F_m_ values are shown for strains grown on TAP. Spot test results and F_v_/F_m_ values are representative of two replicates. **(C)** RT-PCR to detect the expression of the transgene *AtHCF244* in the complemented lines. Primers are listed in Table S1. Actin is used as a loading control. PCR of RNA without reverse transcriptase (-RT) is used to confirm there is no genomic DNA contamination.

### CrHCF244 localizes to chloroplast

Confocal microscopy was used to examine the subcellular localization of CrHCF244. The complementation strain *ami6-C2* (the same line used in Fig. 1A), which is the *ami6* mutant transformed by C2 construct (depicted in Fig. 1C) was used for microscopy. In this strain, CrHCF244 was tagged with fluorescent protein Venus at the C-terminus. As shown in Fig. 6, the Venus signal mostly overlapped with chlorophyll signal, indicating that CrHCF244 is localized in the chloroplast. The strain photosynthetically restored by the Venus-fused CrHCF244 represents the functional and native localization of the protein. Venus tagged CrHCF244 in the background of WT showed the similar localization (Wang *et al*., 2023b).

**Figure 6.**
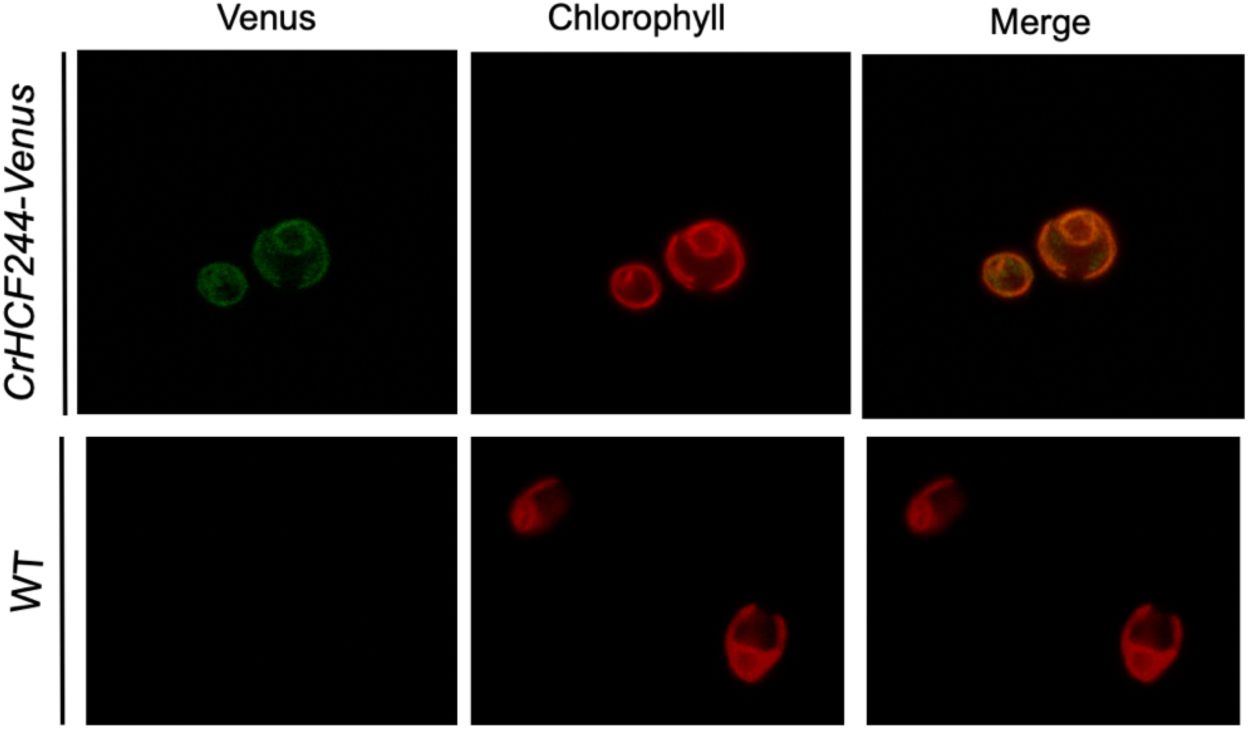
Chloroplast localization of CrHCF244. Venus-tagged CrHCF244 plasmid was transformed into the *ami6* mutant. The complemented line *ami6-C2* was grown under photosynthetic condition before imaging. WT strain was imaged under the same conditions.

### Chlorophyll decreased in the absence of CrHCF244

To test the effect of the absence of *CrHCF244* on the accumulation of chlorophyll, we measured chlorophyll content in the Δ*CrHCF244-T23* mutant grown in the dark and under light, respectively. The total Chl and Chl a/b ratio are shown in Table 1. In the dark, the total Chl decreased around 40% in the mutant and recovered to similar levels of WT in the complemented strains. Under light, the chlorophyll in the mutant also decreased similarly as in the dark. This suggests CrHCF244 could be involved in chlorophyll biosynthesis in *Chlamydomonas*, directly or indirectly through one-helix protein 2 (OHP2), which is involved in chlorophyll binding to apo-D1 and chlorophyll biosynthesis in *Arabidopsis* and *Chlamydomonas* (Hey and Grimm, 2018; Wang *et al*., 2023a). The change of chlorophyll in the Δ*CrHCF244-T23* mutant is similar to *ohp2* mutant of *Chlamydomonas* (Wang *et al*., 2023a).

**Table 1.** Chlorophyll content and Chl a/b ratios in WT, Δ*CrHCF244-T23* and complemented lines. Cells from mixotrophic cultures under light intensity of 35 µmol photons m^-2^ s^-1^ and from heterotrophic cultures in the dark (grey area) in the early exponential phase were used for measurement. Data are presented as means ± standard deviation (n = 3).

| | | [Chl] $\mu\text{g mL}^{-1} \text{OD}_{750}^{-1}$ | Chl-a/b |
| --- | --- | --- | --- |
| light | WT | $33.5 \pm 0.8$ | $3.04 \pm 0.01$ |
| | $\Delta CrHCF244-T23$ | $21.7 \pm 0.6$ | $2.67 \pm 0.05$ |
| | $\Delta CrHCF244-T23-C1$ | $34.6 \pm 1.4$ | $3.08 \pm 0.03$ |
| | $\Delta CrHCF244-T23-C2$ | $33.2 \pm 1.7$ | $3.10 \pm 0.03$ |
| dark | WT | $27.5 \pm 0.4$ | $2.75 \pm 0.17$ |
| | $\Delta CrHCF244-T23$ | $16.9 \pm .07$ | $2.67 \pm 0.14$ |
| | $\Delta CrHCF244-T23-C1$ | $28.0 \pm 2.0$ | $2.71 \pm 0.11$ |
| | $\Delta CrHCF244-T23-C2$ | $28.8 \pm 3.0$ | $2.74 \pm 0.15$ |

### Suppressor mutations suggests an alternative pathway of D1 translational control

Interestingly, we observed colonies originating from Δ*CrHCF244-T23* on minimal medium (HS) plates. Growth and F_v_/F_m_ measurements (Fig. 7A) showed these strains have partially restored PSII activity. PCR and Sanger sequencing of *CrHCF244* proved that the *CrHCF244* gene did not undergo secondary mutations in these colonies (Supplementary Fig. S6A and Fig. S6B), indicating the colonies are likely suppressor mutants that have mutations in other gene(s). This suggests an alternative pathway for D1 translation control. We were also able to observe suppressor mutants in *ami6*, in which 27 genes besides *CrHCF244* have been deleted. We propose the pathway for *psbA* translation control in Fig. S5, in which the alternative pathway is mediated by an alternative translation factor (TF) that was repressed. The identity of the suppressor gene(s) is not investigated here and will be of interest for future study.

**Figure 7.**
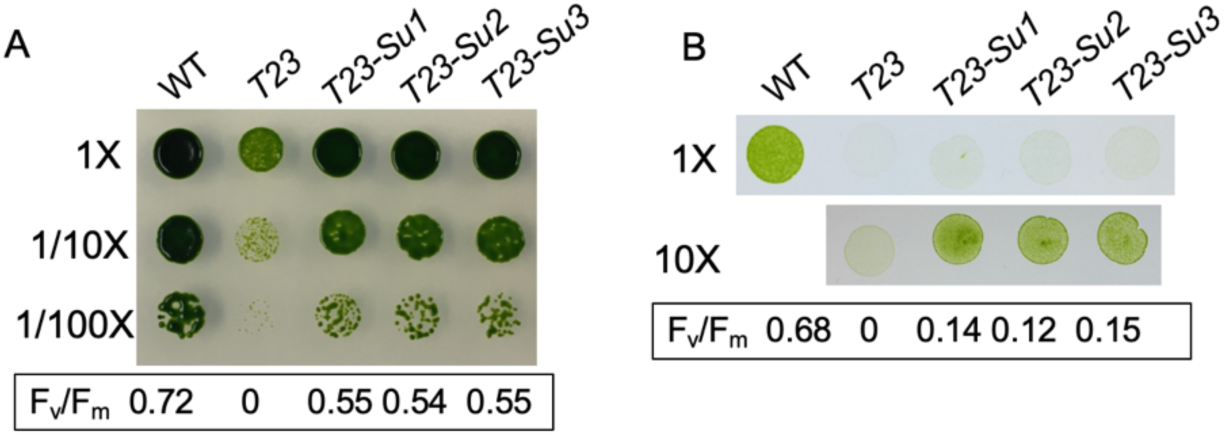
Suppressor mutants of *Δ*CrHCF244-T23 suggest an alternative pathway for D1 translation control. **(A)** Growth and PSII activity of strains under mixotrophic conditions. WT, CRISPR-Cas9 knockout mutant Δ*CrHCF244-T23* (labeled as *T23* here) and three suppressor mutants (labeled as *T23-Su1*, *T23-Su2*, *T23-Su3*). **(B)** Growth and PSII activity of strains, as in (A), under photoautotrophic conditions. Spot test results and F_v_/F_m_ values are representative of two replicates.

### PSII is not affected in CRISPR-Cas9 mutants of RB47 and RB60

To examine the relationship of CrHCF244 with the previously reported D1 translation factors RB47 and RB60, we used CRISPR-Cas9 to generate mutants. The gene maps and Cas9-gRNA RNP cleavage sites are shown in Fig. 8A. We were able to get one mutant for RB47 and multiple mutants for RB60 from PCR screening of the transformants. Here, we used the one mutant of RB47, named as *RB47* and three mutants of RB60, named as Δ*RB60-1*, Δ*RB60-2*, Δ*RB60-3*. PCR identifications of the mutants are shown in Fig. S6A and *RB47* was further verified by Sanger sequencing (Fig. S6B).

**Figure 8.**
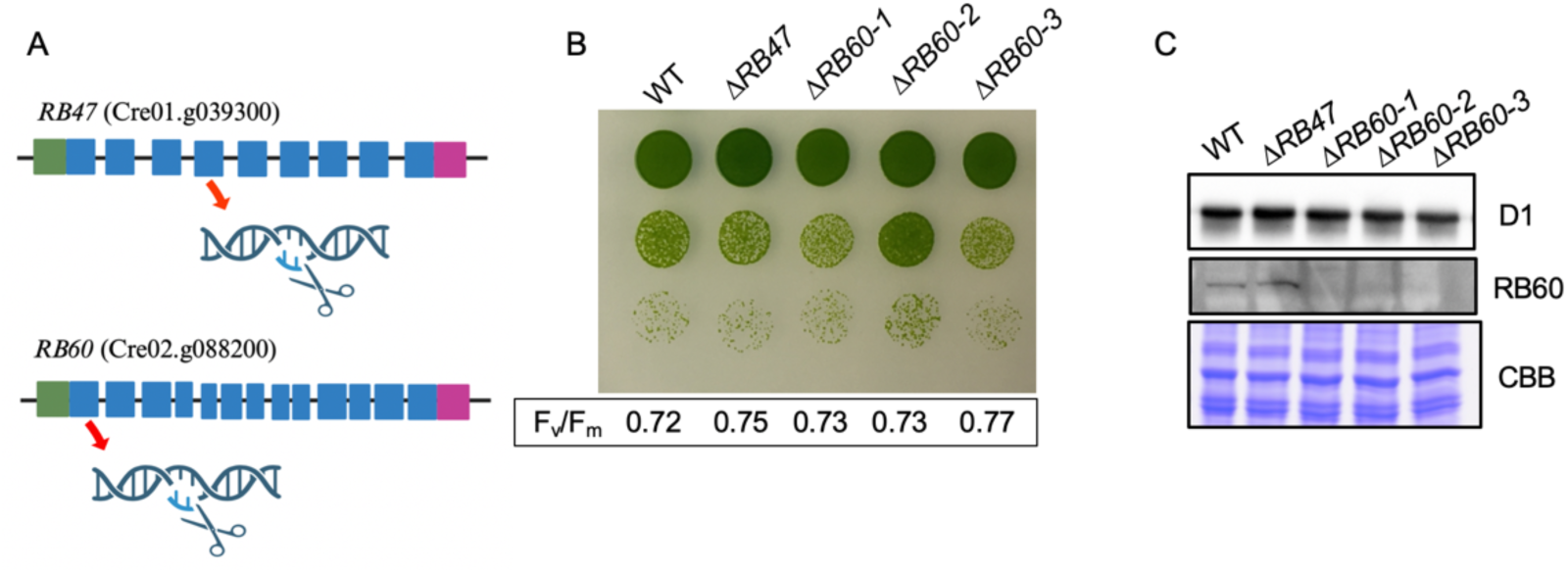
Growth and PSII activity of RB47 and RB60 mutants. **(A)** Gene maps of the *RB47* and *RB60* showing the CRISPR-Cas9 cleavage sites. **(B)** Growth of the *RB47* mutant (*RB47*, see description in text) and RB60 (Δ*RB60-1*, Δ*RB60-2*, Δ*RB60-3*) mutants generated by CRISPR-Cas9, under photoautotrophic conditions. PSII activity was measured as F_v_/F_m_. Spot test results and Fv/Fm values are representative of two replicates. **(C)** Abundance of D1 subunit in the mutants. Immunoblotting of D1 subunit and RB60 was performed in the strains; Coomassie brilliant blue (CBB) stain was shown as a loading control.

*CRISPR-Cas9 editing of RB47* was supposed to introduce a premature stop codon in the fourth exon of the gene (Fig. S6C). Unexpectedly, sequencing of the transcript in the mutant revealed a splicing event that partially removed the insertion (Fig. S6E). The final result is a 23 amino acid insertion. *RB60* was successfully knocked out as evidenced at the DNA and protein levels (Fig. 8C). We note that we were unsuccessful in raising an RB47 antibody.

Unexpectedly, we did not detect significant PSII activity or D1 accumulation defects in the mutants (Fig. 8C). Our genetic analysis does not support the previously proposed roles of RB47 and RB60 in D1 translation (Kim and Mayfield, 1997, 2002; Yohn *et al*., 1998b). This discrepancy may be due to the fact that previous identifications were mostly based on biochemical data when targeted gene editings were not possible.

## Discussion

The high tendency of *Chlamydomonas* to randomly insert DNA fragments into its genome makes it very accessible for generating a mutant pool or library. Such genetic screenings have facilitated the discovery of many protein factors involved in PSII biogenesis and repair including Tba1 (Somanchi *et al*., 2005), REP27 (Park *et al*., 2007), Alb3 (Bellafiore *et al*., 2002; Gohre *et al*., 2006; Ossenbuhl *et al*., 2004), and RBD1 (Calderon *et al*., 2013). Recently, high throughput genetic screening and mapping has resulted in many other potential photosynthesis genes to be identified (Fauser *et al*., 2022; Kafri *et al*., 2023; Li *et al*., 2019a; Wakao *et al*., 2021), which includes the components of photosynthetic apparatus, factors for their biogenesis and maintenance, proteins regulating photosynthesis and other genes whose functions are not yet characterized. In this study, using forward genetics, we identified a mutant, *ami6*, that is defective in PSII (Fig. 1A). To understand PSII biogenesis and identify factors involved in this process, we started by generating a mutant pool and screened it for photosynthetic growth defective mutants using chlorophyll fluorescence to test the PSII activity and found one mutant that does not have PSII activity (Fig. 1A). We then used whole genome sequencing of the mutant and its corresponding WT to map the genomic variation. Analysis of the sequencing data revealed an approximately 345 kb deletion on chromosome 2 (Supplementary Dataset S1) when using the *Chlamydomonas reinhardtii* v5.6 genome (https://phytozome-next.jgi.doe.gov/info/Creinhardtii_v5_6) as the reference genome in early 2022. Recently, we re-analyzed the whole genome sequencing data using *Chlamydomonas reinhardtii* v6.1 (https://phytozome-next.jgi.doe.gov/info/CreinhardtiiCC_4532_v6_1) genome as the reference genome and the deletion (∼347 kb) was mapped onto chromosome 15 (Fig. 1B), due to the significant improvements in assembly quality of the v6.1 genome. Analysis of the whole genome sequencing showed a ∼347 kb deletion, where there are 28 genes, on chromosome 15 in the *ami6* mutant (Fig. 1B). We pinpointed the causative gene to be *Cre02.g142146*, or *CrHCF244*, the homolog gene of *Arabidopsis HCF244* by complementation (Fig. 1A) and by reproducing the mutant phenotype using a CRISPR knockout (Δ*CrHCF244-T23*) of the same gene (Fig. 2).

Both *ami6* and Δ*CrHCF244-T23* have no PSII activity (Fig. 1A and Fig. 2B). Immunoblotting showed that major subunits of PSII are drastically reduced in both mutants, whereas other photosynthetic complexes are not affected (Fig. 3A). The specific effect of CrHCF244 on PSII is consistent with its homologs in plants (Chotewutmontri *et al*., 2020; Link *et al*., 2012). PSII biogenesis is a stepwise and highly coordinated process, and the accumulation of subunits is tightly regulated (Baena-Gonzalez and Aro, 2002; Nickelsen and Rengstl, 2013; Nixon *et al*., 2010). It has been established that the translation of PSII subunits is controlled by an assembly-controlled mode, termed as CES, in which presence of the D2 subunit is prerequisite for translation of D1, and presence of D1 is prerequisite for translation of CP47 (Wollman *et al*., 1999). The mechanism of CES is that the unassembled subunit will bind to the 5’ UTR of its own mRNA and repress its translation. For instance, the translated D1 protein, which hasn’t been assembled into any complex will bind to *psbA* mRNA and repress any more translation of D1 protein. The repression will be released when D1 is assembled into D2-Cyt b_559_ complex (Minai *et al*., 2006). To know whether any defects in synthesis caused the reduced accumulation of PSII subunits, we used ^14^C-acetate pulse labeling and observed that the synthesis of D1 is affected in the mutants (Fig. 4A). As expected, in the absence of D1, CP47 translation was abolished as well in the mutants (Fig. 4A). The data presented showed that CrHCF244 is essential for PSII biogenesis by affecting *psbA* translation, consistent with its role in land plants (Chotewutmontri *et al*., 2020; Link *et al*., 2012).

Due to the large differences in the *psbA* 5’ UTR sequences (Shen *et al*., 2001) and no homolog of trans-regulating factors has been reported to be involved in *psbA* translation between *Chlamydomonas* and *Arabidopsis*, it was not clearly known whether *psbA* translation control is conserved in algae and plants. In this study, we show that *CrHCF244*, a homolog of *Arabidopsis HCF244* function in *psbA* translation, consistent with the function of *HCF244* in *Arabidopsis* (Link *et al*., 2012). The *Arabidopsis* ortholog complementation of the *Chlamydomonas* mutant (Fig. 5B) also demonstrated the functional conservation of this gene from algae to plants, consistent with the relatively high sequence identity (Fig. S7) of the proteins between the two organisms. The other *psbA* translation factor identified in *Arabidopsis*, HCF173 (Schult *et al*., 2007), is also possibly involved in *psbA* translation in *Chlamydomonas*. In a proteomics study, it was noted that D1 accumulation was reduced in a *CrHCF173* mutant (Kafri *et al*., 2023) which suggests it could be a translation factor for *psbA*. Previously RB47 and RB60, were biochemically identified as possible *psbA* translation factors in *Chlamydomonas*. However, genetic evidence in this study showed that RB47 and RB60 play no major role in *psbA* translation (Fig. 8B, 8C, 8D). No *Arabidopsis* homologs of these proteins functioning in *psbA* translation have been identified. These data also support the hypothesis that similar factors are being employed in *psbA* translation in the two organisms.

We observed less chlorophyll content in Δ*CrHCF244-T23* compared to the WT and complemented lines (Table 1), which resembles the result found in the *Chlamydomonas ohp2* mutant (Wang *et al*., 2023a). In plants, OHP1 and OHP2 bind and deliver chlorophyll to apo-D1. OHP1, OHP2, and HCF244 form a complex and there is interdependence of protein accumulation among the components (Chotewutmontri and Barkan, 2020; Hey and Grimm, 2018; Li *et al*., 2019b; Myouga *et al*., 2018). Future investigation will explore the relationship between CrHCF244, OHP2, and chlorophyll biosynthesis in *Chlamydomonas*.

Previously, it was proposed that RB60 regulates the redox state of RB47, promoting its binding to *psbA* mRNA and translation (Kim and Mayfield, 1997). Here we showed that neither of these proteins is essential for *psbA* translation using genetic evidence (Fig. 8). The previous conclusions were mostly based on *in vitro* biochemical data (Danon and Mayfield, 1991; Kim and Mayfield, 1997; Yohn *et al*., 1998a). There was no genetic study on RB60, and the genetic analysis on RB47 used mutants with reduced accumulation of RB47 but no mutation in the *RB47* gene (Bruick and Mayfield, 1999; Yohn *et al*., 1998b), due to the limitation in targeted mutagenesis of nuclear genes in *Chlamydomonas*. A previous scenario of unspecific binding of *psbA* mRNA was observed in which RB38, originally identified as a *psbA* mRNA associating protein (Danon and Mayfield, 1991), was later shown to function in *psbD* translation control (Schwarz *et al*., 2007).

In this study, we confirmed that CrHCF244 is a *psbA* translation factor. A previous study identified Tba1 as a *psbA* translation factor (Somanchi *et al*., 2005) in *Chlamydomonas*. There was a suggestion that CrHCF173 is also a *psbA* translation factor based on the function of its *Arabidopsis* homology and proteomics detection of reduced D1 accumulation in the *Chlamydomonas* mutant (Kafri *et al*., 2023). Therefore, there are at least two or three proteins required for translation of *psbA in Chlamydomonas*, considering DLA2 is a moonlighting regulator, which only functions in the presence of acetate (Bohne *et al*., 2013; Neusius *et al*., 2022). Questions arise as to why multiple components are required for the translation of one message and how the absence of each of them would abolish the translation. They could function in a complex with each of them required for the stability of the complex, like the translation initiation complex in the cytosol of eukaryotes (Brito Querido *et al*., 2024; Jackson *et al*., 2010). Alternatively, the translation initiation of *psbA* could be a sequential process and each of them is needed in a stepwise way. Regarding the working mechanism of CrHCF244 in *psbA* translation, this protein could directly bind to the 5’ UTR of *psbA* or bind indirectly as mediated by other proteins. There are proteins in the SDR superfamily shown to be RNA binding proteins (Bollenbach and Stern, 2003). We tried to recombinantly express CrHCF244 and test its RNA binding ability *in vitro* but failed to get soluble expression of the protein in *E. coli*.

Our observation of suppressor mutants of Δ*CrHCF244* suggests the presence of an alternative complex for D1 translation or perhaps a step in the translation process could be bypassed in these lines. The inability of the Δ*CrHCF244-T23* mutant to grow photoautotrophically and the absence of PSII activity indicate this alternative pathway is in repression. We propose a possible scenario as illustrated in Fig. S5. Another possibility is mutation in the 5’ UTR of the target gene in the background of a translation factor mutant would (partially) restore the phenotype. The mutation in the 5’ UTR led to a secondary structure change which allowed access for ribosome binding in the absence of the translation factor. A suppressor mutation of a translation factor of *psbD* was reported and the mutation is in the 5’ UTR of the *psbD* mRNA (Rochaix *et al*., 1989). We sequenced the *psbA* 5’ UTR of these suppressors and there is no mutation, suggesting the mutation might involve nuclear gene product(s).

This study shows that CrHCF244 is essential for PSII biogenesis by affecting *psbA* translation. This identification of a plant homolog involved in *psbA* translation demonstrates the conservation of *psbA* translation control between algae and plants. Previously, it had been speculated that *psbA* translation control might be distinct between algae and plants due to the large difference in the 5’ UTR of *psbA* genes, and no shared translation factor proteins had been identified between the organisms. The establishment of the conservation of *psbA* translation control will allow the use of *Chlamydomonas*, which has more feasible genetic and systematic mutant screen tools to facilitate the understanding of this important process in plants. The observation of partial suppressors of *CrHCF244* mutants suggests an alternative pathway in *psbA* translation control. Future mapping of the mutation gene(s) in the suppressors and studies of the plant homolog(s) will improve our understanding in *psbA* translation control.

## Abbreviations

PSII: Photosystem II
ROGE: regulators of organelle genes
HCF244: high chlorophyll fluorescence 244
At: *Arabidopsis thaliana*
Cr: *Chlamydomonas reinhardtii*
cTP: chloroplast transit peptide
cpPDC: chloroplast pyruvate dehydrogenase
DLA2: dihydrolipoamide acetyltransferase subunit 2
BN-PAGE: Blue native polyacrylamide gel electrophoresis
TF: translation factor

## Supplementary data

Supplementary data for this manuscript is listed as below and available online.

*Figure S1.* The large deletion in *ami6* was verified by PCR and Sanger sequencing.

*Figure S2.* Genomic identification of Δ*CrHCF244-T23* mutant.

*Figure S3.* BN-PAGE separation of photosynthetic protein complexes.

*Figure S4.* PCR and Sanger sequencing verification *CrHCF244* in the suppressor mutants of Δ*CrHCF244-T23*.

*Figure S5.* Proposed working model of alternative *psbA* translation control.

*Figure S6.* Genomic identification of CRISPR-Ca9 mutants of *RB47* and *RB60*.

*Figure S7.* Clustal Omega multiple sequence alignment of HCF244 from *Chlamydomonas* (CrHCF244) and *Arabidopsis* (AtHCF244).

*Table S1.* Primers and crRNA sequences used in this study.

*Dataset S1. Genomic variance and deleted gene list in ami6*

## Acknowledgments

We thank Shared Instrumentation Facility at Louisiana State University for confocal microscopy support. We thank Prof. Alice Barkan for helpful discussions. We thank Prof. Jun Minagawa for the gift of the B-Hiss *Chlamydomonas* strain. We thank Prof. Terry Bricker for the Cyt f antibody. We thank Dr. Yuqing Hou at University of Massachusetts Medical School for her advice on CRISPR-Cas9 gene editing in *Chlamydomonas*. BioRender was used for making diagrams (Fig. 1C, Fig.2A, Fig. 5A, Fig. S5 and Fig. 8A)

## Author contributions

Conceptualization: XW, JVM, DJV; Experiments: XW, LAC, CZ, JVM, DJV; Data analysis and interpretation: XW, GW, CZ, MD, JVM, DJV; Writing: original draft-XW, JVM, DJV, reviewing and editing-all authors; Supervision of study: DJV; Funding acquisition: DJV and MD.

## Data availability

All relevant *Chlamydomonas* strains will be archived in Chlamydomonas Resource Center. All relevant data and other materials are available from the authors upon request.

