## Supporting Information for "*CrHCF244* is required for *psbA* translation in *Chlamydomonas reinhardtii*"

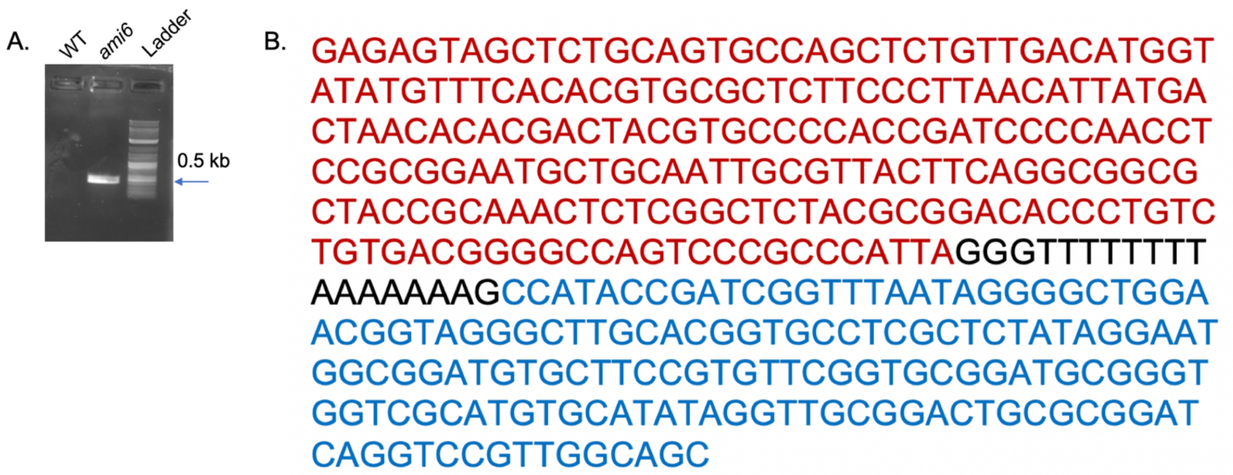


***Figure S1. The large deletion in ami6 was verified by PCR and Sanger sequencing.***

**(A)** PCR identification of *ami6* genetic background flanking the large deletion, using primers in Table S1. A band with size of around 0.5 kb was amplified from *ami6*, whereas no PCR product was amplified in WT. **(B)** Sanger sequencing and alignment result of *ami6* PCR product in A. The sequence in red is part of Cre02.g141766, the sequence in blue is the intergenomic sequence between Cre15.g801755 and Cre15.g801756 and the sequence in black is unknown sequence, which could be a small insertion when the large deletion occurred.

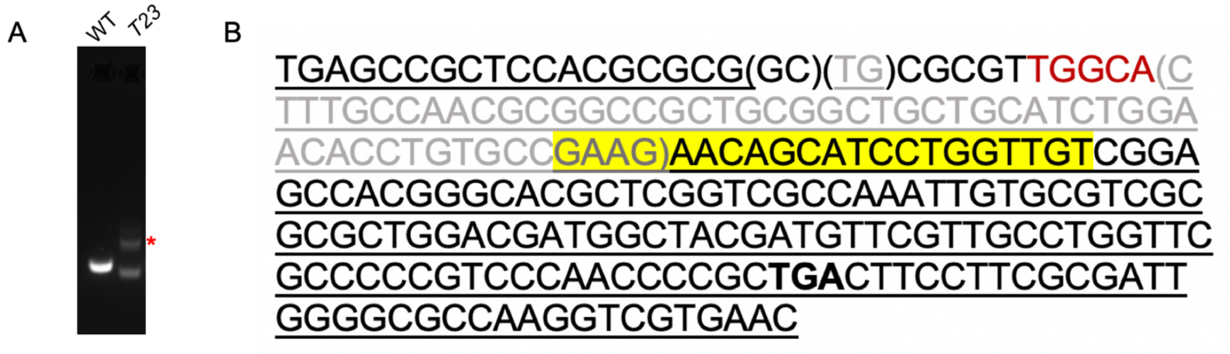


***Figure S2. Genomic identification of ∆CrHCF244-T23 mutant.***

**(A)** PCR identification showed a small deletion in ∆*CrHCF244-T23* (shown as *T23* in the figure). Red asterisk indicates the unspecific band. **(B)** Sequence of exon 2 of *CrHCF244* in ∆*CrHCF244-T23* by Sanger sequencing. Underlines are original sequence in WT genome. GC in parentheses is a replacement of TG. Red is the inserted 5 nucleotides and grey in parentheses is the 52 nucleotides being deleted. The insertion and deletion created a premature stop codon as shown in bold and underline.  Yellow highlight is the guide RNA (gRNA) binding site.

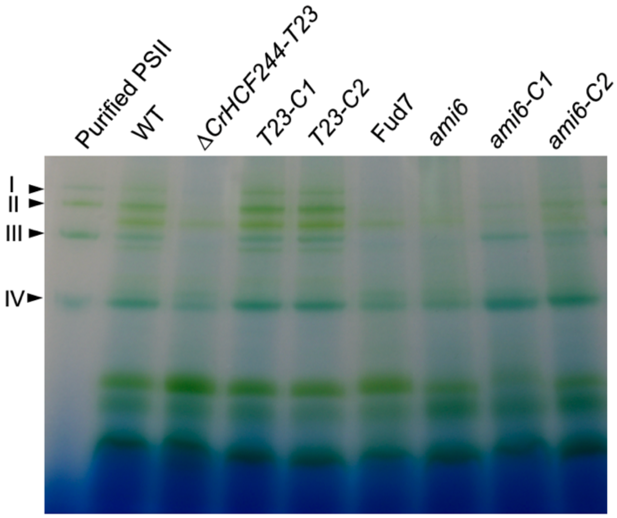


***Figure S3. BN-PAGE separation of photosynthetic protein complexes.***

Lane 1 is purified PSII complexes from B-His strain which has 6 His tag on the C-terminal of PsbB. Arrows indicate the positions of different PSII complexes. Lanes 2-9 are solubilized thylakoid membrane from strains as indicated.

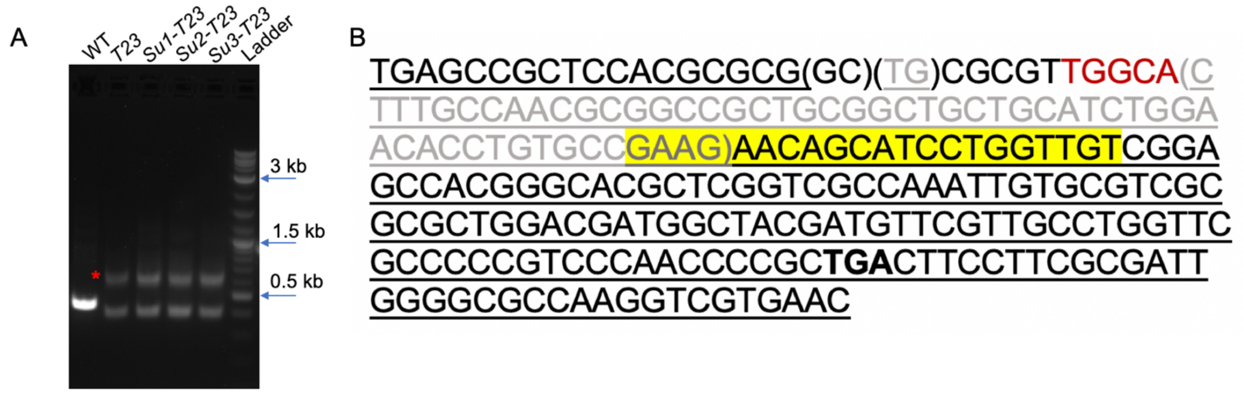


***Figure S4. All three strains gave the same sequencing, which is identical with the original mutant strain, showing the interrupted CrHCF244 gene is not restored in those suppressors.***

**(A)** PCR identification of the three suppressors using the same primers for PCR identification Δ*CrHCF244-T23* (shown as T23 in the figure) in Table S1. Red asterisk indicates the unspecific band. **(B)** Sequence of exon 2 of *CrHCF244* in the three suppressors by Sanger sequencing. Underlines are original sequence in WT genome. GC in parentheses is a replacement of TG. Red is the inserted 5 nucleotides and grey in parentheses are the 52 nucleotides being deleted. The insertion and deletion created a premature stop codon as shown in bold and underline. Yellow highlight is the guide RNA (gRNA) binding site.

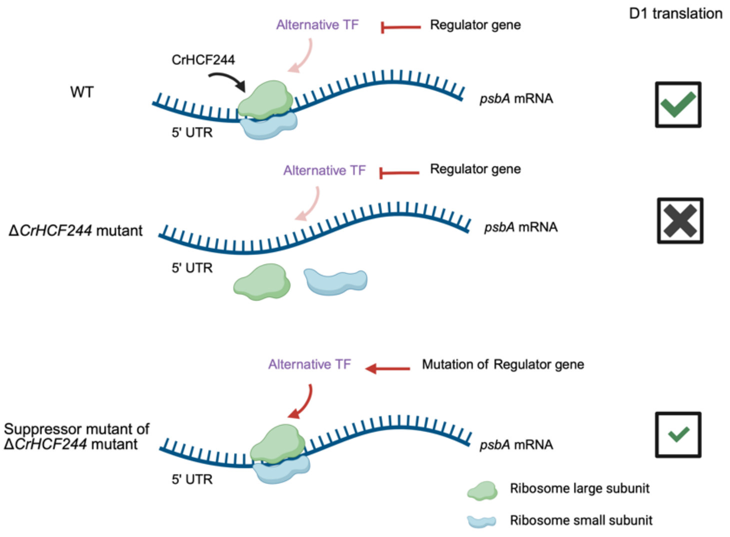


***Figure S5. Proposed working model of alternative psbA translation control.***

In the absence of CrHCF244, the ribosome cannot bind *psbA* mRN thus there is no D1 translation in the Δ*CrHCF244-T23* mutant. A mutation in a regulator gene activates an alternative translation factor (TF), which allows ribosome loading to *psbA* mRNA, thus enabling D1 translation in the suppressor mutants.

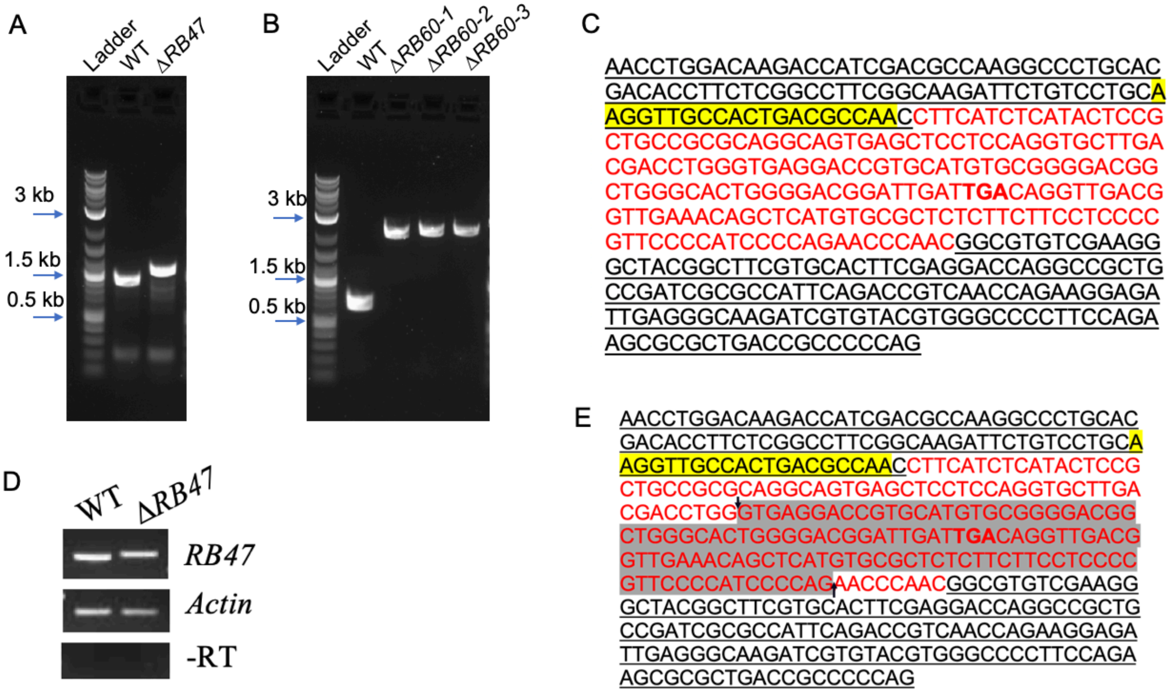


***Figure S6. Genomic identification of CRISPR-Ca9 mutants of RB47 and RB60.***

**(A)** PCR identification of Δ*RB47* using primers in Table S1. Primers flanking Cas9 cleavage site. **(B)** PCR identification of Δ*RB60* mutants using primers in Table S1. Primers flanking Cas9 cleavage site. **(C)** Sequence of exon 4 of *RB47* in ΔRB47 by Sanger sequencing. Underlines are original sequence in WT genome; red is 184 nucleotides insertion with the resulting premature stop codon in red bold; yellow highlight is the guide RNA (gRNA) binding sequence. **(D)** RT-PCR of *RB47* transcription in Δ*RB47* using primers in Table S1. The size difference between WT and Δ*RB47* is due to an insertion, as shown in **(E)**. Actin is used as a loading control. Treatment of RNA without reverse transcriptase (bottom panel) is used to verify there is no genomic DNA contamination. **(E)** Sanger sequencing of the PCR product in upper panel of **(D)**. Underlines are original sequence in WT; yellow highlight is the guide RNA (gRNA); red is 69 nucleotides insertion; the shaded area of 115 bp which is in the DNA sequence of Δ*RB47* is spliced out, possibly using the GT and AG as splicing sites as indicated by the arrows.

```
AtHCF244      MASLRLPAQLVTRGNLIHHNSSSSSSGRLSWRRSLTPENTIPLFPSSSSSSLNRERSIVV      60
CrHCF244      -----------------MQTANSVAGQRLALRQG-----VVAFLPVRAP----VSRSTRV      34
                                :.:.* :. **: *:.     .: ::*  :      .**  *

AtHCF244      PVTCSAAAVNLAPGTPVRPTSILVVGATGTLGRQIVRRALDEGYDVRCLVRPRPAPADFL      120
CrHCF244      RVFANAAAAAAASGTPVPKNSILVVGATGTLGRQIVRRALDDGYDVRCLVRPRPNPADFL      94
               * ..***.  * ****  .*********************:************ *****

AtHCF244      RDWGATVVNADLSKPETIPATLVGIHTVIDCATGRPEEPIKTVDWEGKVALIQCAKAMGI      180
CrHCF244      RDWGAKVVNGDLTDPSSIPACLVGVNAVIDCATARPEESTRKVDWEGKVALIQSAQAMGI      154
              *****.***.**:.*.:*** ***::******.****  :.************.*:****

AtHCF244      QKYVFYSIHNCDKHPEVPLMEIKYCTEKFLQESGLNHITIRLCGFMQGLIGQYAVPILEE      240
CrHCF244      QRYVFFSIFDCDKHPQVPLMNIKSCTEKFLEASSLDYTTFRLCGFHQAVIGNYAVPILEE      214
              *:***:**.:*****:****:** ******: *.*:: *:***** *.:**:********

AtHCF244      KSVWGTDAPTRVAYMDTQDIARLTLIALRNEKINGKLLTFAGPRAWTTQEVITLCERLAG      300
CrHCF244      RSVWGTNDETRTAYLDSQDVAKMTMAALRGDKTSRKTLPLSGPKAWTTKEVIELCEKMAD      274
              :*****:  **.**:*:**:*::*: ***.:* . * * ::**:****:*** ***::*.

AtHCF244      QDANVTTVPVSVLRVTRQLTRFFQWTNDVADRLAFSEVLSSDTVFSA---PMTETNSLLG      357
CrHCF244      TNAKVTTVPTWLLKRTRGVLRSMQWAADAADRLAFAEVLSNNEVWQASAADMAETYRLLD      334
               :*:*****. :*: ** : * :**: *.******:****.: *:.*    *:**  **.

AtHCF244      VDQKDMVTLEKYLQDYFSNILKKLKDLKAQSKQSDIYF    395
CrHCF244      MDPGSVTDLETYLQEYFSRILKKLKEVGASADRTNFYV    372
              :*  .:. **.***:***.******:: *.:.::::*.
```

***Figure S7. Clustal Omega multiple sequence alignment of HCF244 from Chlamydomonas (CrHCF244) and Arabidopsis (AtHCF244).***

The alignment showed 56.92% identity.

***Supplementary Table S1. Primers and crRNA sequences used in this study.***

| ***Primer name*** | ***Sequence*** | ***Purpose*** |
|---|---|---|
| ami6_1 | CAGCTACCCGCCCGATGA | Verification of the large deletion in *ami6* by Sanger sequencing |
| ami6_2 | GCTGCCAACGGACCTGATCC | |
| CGL102_start | ATGCAGACAGCAAACAGC | Amplifying ORF of *CrHCF244* |
| CGL102_stop | TTACACGTAGAAGTTGGTGCG | |
| CGL102_F_ pLM006 | GCTACTCACAACAAGCCCAGTTATGCAGACA GCAAACAGC | PCR for linear vector to construct C1 plasmid |
| CGL102_R_ pLM006 | CCTGAGAATTCGTGCAGTTATTACACGTAGAA GTTGGTGCGGTC | |
| CGL102_F_ pLM006 | GCTACTCACAACAAGCCCAGTTATGCAGA CAGCAAACAGC | PCR for linear vector to construct C2 plasmid |
| cgl102_pLM005 fusion | GAGCCACCCAGATCTCCGTTCACGTAGA AGTTGGTGCG | |
| AtHCF244_F | GCTACTCACAACAAGCCCAGTT | Amplifying CrPsaD-mAtHCF244 |
| AtHCF244_R | CCTGAGAATTCGTGCAGTTA | |
| pLM006_F_no tag | taactgcacgaattctCAGG | PCR for linear vector to construct AtHCF244 plasmid |
| pLM006_R | AACTGGGCTTGTTGTGAGTAGC | |
| crRNA for *CrHCF244* | 5'-/AltR1/rArCrA rArCrC rArGrG rArUrG rCrUrG rUrUrC rUrUrG rUrUrU rUrArG rArGrC rUrArU rGrCrU /AltR2/ -3' | crRNA for CRISPR-Cas9 editing *CrHCF244* |
| cgl102-40HR_F | TGCCAACGCGGCCGCTGCGGCTGCTGCATCT GGAACACCTGTAAAACGACGGCCAGTGAG | Donor DNA for CRISPR-Cas9 editing *CrHCF244* |
| cgl102_40HR_R | GCGCGCGACGCACAATTTGGCGACCGAGCGT GCCCGTGGCCTGGCACGACAGGTTTCCCG | |
| CGL102_start | ATGCAGACAGCAAACAGC | PCR screening of Δ*CrHCF244* transformants |
| CGL102_E4R | CTGGGGATGTTTGTCGCA | |
| crRNA for *RB47* | 5'-/AltR1/rArArG rGrUrU rGrCrC rArCrU rGrArC rGrCrC rArArG rUrUrU rUrArG rArGrC rUrArU rGrCrU /AltR2/ -3' | crRNA for CRISPR-Cas9 editing *RB47* |
| RB47_40HR_F | CCCTGCACGACACCTTCTCGGCCTTCGGCAAGATTC TGTCGAATTCGATATCAAGCTTCT | Donor DNA for CRISPR-Cas9 editing *RB47* |
| RB47_40HR_R | GGCCTGGTCCTCGAAGTGCACGAAGCCGTAGCCC TTCGACCGCTTCAAATACGCCCAG | |

| ***Primer name*** | ***Sequence*** | ***Purpose*** |
|---|---|---|
| RB47_I3_F | CAGCAGTGACATGACCTG | PCR screening of Δ*RB47* transformants |
| RB47_I5_R | TGTGAATGCTCCCGTGTG | |
| crRNA for *RB60* | 5'- /AltR1/rCrUrG rUrGrG rUrGrA rCrCrG rUrCrA rArGrA rArCrGrUrUrU rUrArG rArGrC rUrArU rGrCrU /AltR2/ -3' | crRNA for CRISPR-Cas9 editing *RB60* |
| RB60_40HR_F | GATGCCCCCGCCGCCCCTAAGGACGACGACGTCGA CGTTAGAATTCGATATCAAGCTTCT | Donor DNA for CRISPR-Cas9 editing *RB60* |
| RB60_40HR_R | CTCACTCCACAAGCGCGAACTTGGACTTCTTGACGG TCTCCGCTTCAAATACGCCCAG | |
| RB60_start | ATGAACCGTTGGAACCTTC | PCR screening of Δ*RB60* transformants |
| RB60_E4_R | CGCCATCAACGAACCACTTG | |
| psbA_E2E3_F | CTACATGGGTCGTGAGTGGG | qPCR of *psbA* |
| psbA_E3R | ATGAACCTTGGCCGATAGGG | |
| psbA_E5F | GCTTGGCCGGTAATCGGTAT | qPCR of *psbA* |
| psbA_E5R | CGGTTGATGATGTCTGCCCA | |
| psbB_F | GTGTTGCAGCAGCTCACATT | qPCR of *psbB* |
| psbB_R | ACGTGGGTCACGGAAAAGTT | |
| psbD_F | GTTTGGGGTCCAGAAGCTCA | qPCR of *psbD* |
| psbD_R | ATGCACCGTGTAAAGCAACG | |
| Actin_F | GCCAGAAGGACTCGTACGTT | RT-PCR of *Actin* |
| Actin_R | CGCCAGAGTCCAGCACGATA | |

| | | |
|---|---|---|
| RB47E4_F | GGCCTTCGGCAAGATTCTGT | RT-PCR of *RB47* |
| RB47E7_R | AGAGTTGGCGAACAGCTCAC | |
| AtHCF244_1F | TCGTGAACGCTGATCTCTCG | RT-PCR of *AtHCF244* |
| AtHCF244_2R | GGTACTTCTCCAGCGTCACC | |
